# New targets acquired: improving locus recovery from the Angiosperms353 probe set

**DOI:** 10.1101/2020.10.04.325571

**Authors:** Todd G.B. McLay, Joanne L. Birch, Bee F. Gunn, Weixuan Ning, Jennifer A. Tate, Lars Nauheimer, Elizabeth M. Joyce, Lalita Simpson, Nick Weigner, Alexander N. Schmidt-Lebuhn, William J. Baker, Félix Forest, Chris J. Jackson

## Abstract

Universal target enrichment kits maximise utility across wide evolutionary breadth while minimising the number of baits required to create a cost-efficient kit. Locus assembly requires a target reference, but the taxonomic breadth of the kit means that target references files can be phylogenetically sparse. The Angiosperms353 kit has been successfully used to capture loci throughout angiosperms but includes sequence information from 6–18 taxa per locus. Consequently, reads sequenced from on-target DNA molecules may fail to map to references, resulting in fewer on-target reads for assembly, reducing locus recovery. We expanded the Angiosperms353 target file, incorporating sequences from 566 transcriptomes to produce a ‘mega353’ target file, with each gene represented by 17–373 taxa. This mega353 file is a drop-in replacement for the original Angiosperms353 file in HybPiper analyses. We provide tools to subsample the file based on user-selected taxon groups, and to incorporate other transcriptome or protein-coding gene datasets. Compared to the default Angiosperms353 file, the mega353 file increased the percentage of on-target reads by an average of 31%, increased loci recovery at 75% length by 61.9%, and increased the total length of the concatenated loci by 30%. The mega353 file and associated scripts are available at: https://github.com/chrisjackson-pellicle/NewTargets

## INTRODUCTION

Target enrichment (also known as target capture, exon capture, HybSeq) has become the leading high-throughput sequencing methodology for phylogenomics, offering reliable retrieval of hundreds or thousands of loci at a reasonable price per base pair (bp) (Cronn et al., 2012; Grover et al., 2012; Barrett et al., 2016; Bragg et al., 2016). The method has proven useful for resolving relationships at all taxonomic scales, including higher level phylogenetic relationships between orders or families, as well as lower level relationships between genera or species, and for species delimitation (Bi et al., 2013; Nicholls et al., 2015; Song et al., 2017; Choi et al., 2019; Breinholt et al., 2019). Target enrichment uses available genome sequence information in the form of genomes, transcriptomes, or genome skimming data in order to identify a set of target loci (e.g. genes, exons, or Ultra Conserved Elements (UCEs)), that are typically low or single copy (Faircloth 2017; McKain et al., 2018). From the target loci set, short 80–120 bp RNA baits (also called probes) are designed, to create a ‘bait kit’. These short RNA baits are used in a hybridisation reaction to bind to DNA fragments matching the target loci, which are then captured and PCR-amplified for sequencing. The increasing availability of genomic resources held in public repositories, combined with pipelines to identify low-or-single-copy genes based on these resources, have enabled bait kit design for a wide range of plant groups (Kadlec et al., 2017; Campana et al., 2018; Chafin et al., 2018; Vatanaprast et al., 2018).

Universal bait kits, such as the Angiosperms353 bait kit, aim to capture the same loci set from samples representing significant phylogenetic breadth and evolutionary time (Bossert et al., 2018; Breinholt et al., 2019; Johnson et al., 2019). Such kits typically require a larger number of baits to encompass the sequence diversity potentially found between samples at each locus. Larger kits are more costly (Hutter et al., 2019; Couvreur et al., 2019), and therefore to keep costs manageable universal bait kits balance the number of baits synthesised, and hence bait sequence diversity for each locus, against the total number of RNA baits strictly required to fully capture diversity at each locus. Incomplete representation of sample sequence diversity in the synthesised baits is in part compensated for by the high affinity of the biochemical interaction in the hybridisation reaction binding the RNA-bait to the DNA-target. This high affinity means that target DNA can be successfully captured even in cases where bait and target sequences differ by ∼20% (though Johnson et al., 2019 extended this to 30% when designing the Angiosperms353 kit) and provides a constraint around the minimal sequence diversity required to capture loci across the desired phylogenetic breadth (Mayer et al., 2016; Branstetter et al., 2017; Faircloth 2017; Couvreur et al., 2019). This is demonstrated by the wide range of flowering plant groups that have successfully utilised the Angiosperms353 kit (Johnson et al., 2019; Van Andel et al., 2019; Larridon et al., 2020; Shee et al., 2020) as well as many other universal bait kits e.g. flagellate plants – ‘GoFlag’ (Breinholt et al., 2019); ferns (Wolf et al., 2018); arachnids (Starrett et al., 2016); Cnidaria (Quattrini et al., 2018); and Gastropoda (Teasdale et al., 2016).

Assembly of raw sequence reads into the desired locus typically follows one of two strategies; 1) de-novo assembly of reads and subsequent matching of contigs to targets loci, or 2) mapping reads to each locus, followed by de-novo assembly of the mapped reads for each locus. Various pipelines are available to perform locus assembly, such as HybPiper (read-mapping; Johnson et al., 2016), PHYLUCE (de-novo assembly; Faircloth 2016), and SECAPR (both de-novo and read-mapping possible; Andermann et al., 2018). For either strategy, a file containing the loci targeted (i.e. the target file) is required. This is typically the same file that was used to design the baits. For universal-scale kits this means that closely related reference sequences might not be present in the target file for a given dataset. This raises a question: what if the biochemistry of hybrid-enrichment enables the successful capture of target loci DNA *in vitro*, but subsequent bioinformatic processing of raw or assembled data to reconstruct the target locus is inefficient or fails because there is no suitable reference *in silico*? A mismatch between biochemical locus capture and bioinformatic locus recovery will have a larger impact in broader-scale universal kits, or groups where suitable reference sequences are lacking, and could influence locus recovery at any phylogenetic level. To investigate the impact of target file sequence diversity on locus recovery we developed tools to expand the Angiosperms353 target file and compared locus recovery across a range of phylogenetic depths against the default 353 file, using HybPiper (Johnson et al., 2016) for locus assembly.

## METHODS AND RESULTS

### Generating the mega353 target file

The target file for the Angiosperms353 kit was downloaded from https://github.com/mossmatters/Angiosperms353/blob/master/Angiosperms353_targetSequences.fasta, here-on referred to as the ‘default353’ target file. To obtain a phylogenetically diverse set of angiosperm sequences from which to recover the Angiosperms353 loci, transcriptomes were downloaded from the 1KP portal (http://www.onekp.com/public_data.html; Carpenter et al., 2019). A maximum of two samples per genus were added, with samples with the largest number of sequences preferentially included (see https://github.com/chrisjackson-pellicle/NewTargets - ‘control file’). The resulting set included 566 transcriptomes.

To create the mega353 target file, the following process was carried out (summarised in Fig. 1). For each gene in the default353 target file a single gene alignment was produced using MAFFT (Katoh and Standley 2013), and a corresponding Hidden Markov Model (HMM) profile was generated using HMMER (Eddy 2011). HMM profiles were used to search the 1KP transcriptomes using hmmsearch with an eValue cut-off of 1e-50, and the top hit (if present) was recovered. Transcriptome hits were added to the corresponding gene alignment, and the 5’ and 3’ termini were trimmed to the longest original target file sequence from either *Arabidopsis thaliana* (L.) Heynh., *Amborella trichopoda* Baill., or *Oryza sativa* L., as at least one of these three species was included for each locus in the default353 target file. In cases where a transcriptome hit sequence was <85% the length of the longest original target file sequence for a given gene, the closest related target file sequence was identified using a distance matrix, and the transcriptome hit sequence was extended by grafting with the 5’ and/or 3’ termini of the closest related sequence. The resulting target file sequence was therefore a chimeric construct, and these cases are flagged in the sequence name. This grafting process was necessary as HybPiper translates a single chosen target file sequence for each gene and sample, and the resulting protein sequence is used as a query in Exonerate (Slater and Birney 2005) to search against assembled nucleotide contigs, using the protein2genome model. Consequently, short protein queries recover truncated nucleotide loci sequences, even if longer contigs have been successfully assembled.

**Figure 1:**
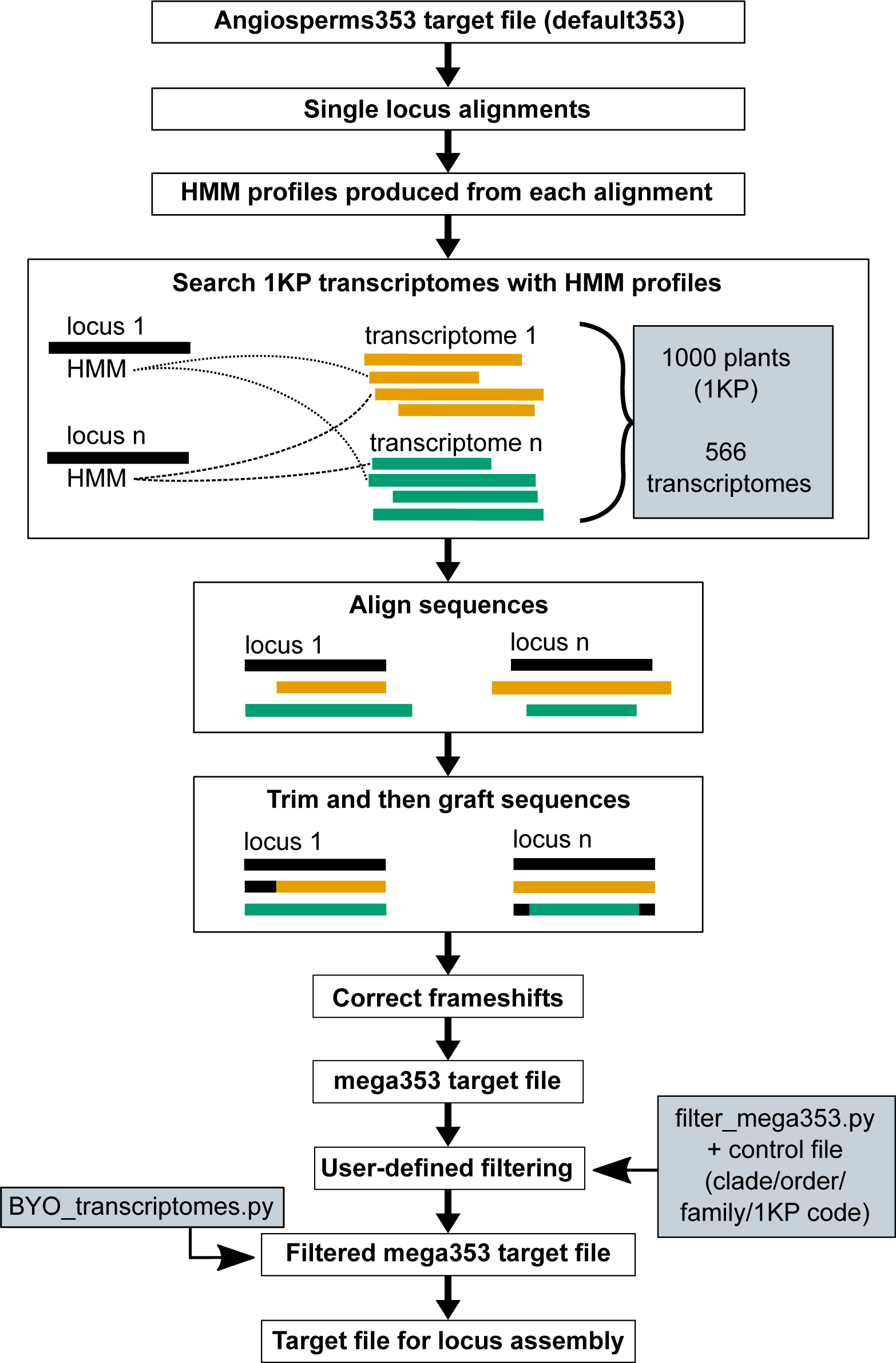
Overview of the steps involved in creating the mega353 target file. Firstly, loci in the default353 file are aligned and HMM profiles are created for each locus. The HMM profiles are used to identify those loci in the 1 KP transcriptomes (ts), which are added to the alignment. The alignment of each locus is then trimmed and grafted, and a frameshift correction is performed, and all loci are combined in the mega353 target file. The mega353 target file can then be filtered using the control file to set which samples to be included in the final target file for locus assembly. The BYO_transcriptomes.py script can be used to add GenBank or personal transcriptomes to the filtered mega353 target file.

As recovery of target loci using HybPiper requires correct translation of chosen target file sequences in the first reading frame, any frameshifts observed in trimmed and/or grafted transcriptome hit sequences were corrected or compensated for (see https://github.com/chrisjackson-pellicle/NewTargets for further details). In cases where a frameshift could not be corrected, the corresponding transcriptome hit sequence was removed for that gene/sample. Finally, sequences were extracted from each gene alignment, gap positions were removed, and all sequences were concatenated to create a new target file.

In the default353 file there are 4780 target reference sequences and each gene is represented on average by 13.5 reference sequences (range 6–18). In the mega353 target file there are 98,994 target reference sequences and each gene is represented by an average 280 reference sequences (range 17–373). In terms of improvement in phylogenetic density, the default353 target file has an average of 13.5 orders and 13.5 families per gene, whereas the mega353 target file has an average of 49.8 orders and 170 families per gene (Fig. 2, Supplementary Fig. 1, Supplementary Table 1).

**Figure 2:**
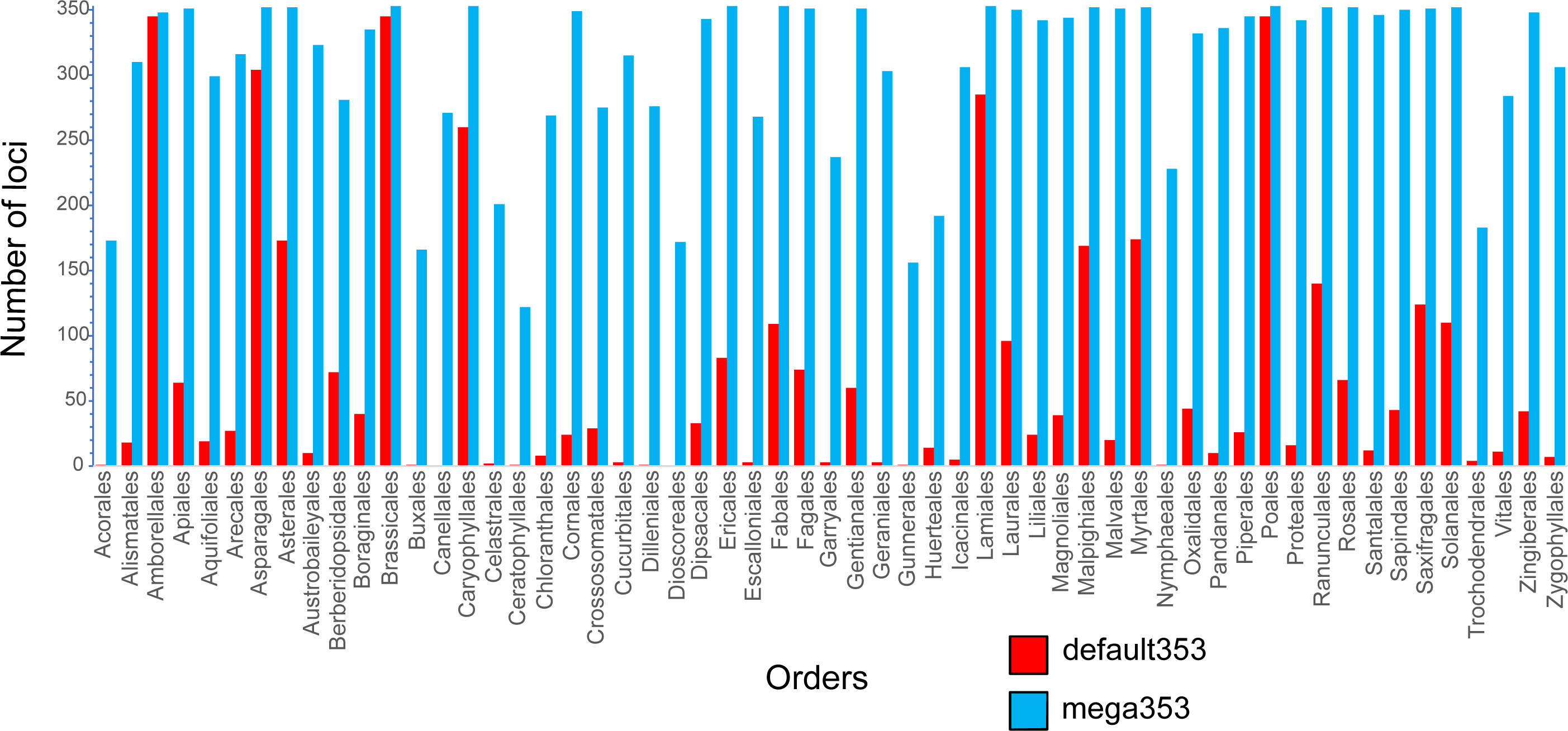
Comparing the number of genes represented for each order in the default353 (red) or mega353 (blue) target files.

### Filtering the mega353 target file

To tailor the large mega353 target file to investigation-specific taxon sampling, we include the script filter_megatarget.py. This script can be used to create a filtered target file based on user-selected taxa or taxon groups, defined by unique 1KP transcriptome codes, families, orders, or clades (see https://github.com/chrisjackson-pellicle/NewTargets for full options). In addition to the chosen samples, all sequences from the default353 target file are retained.

### Adding sequences from any transcriptome to any existing target file

As an additional resource, we provide the script BYO_transcriptomes.py, allowing sequences from any transcriptome (e.g. from GenBank or personal data) to be added to an existing target file. A target file and a directory of transcriptomes are the only inputs required. For Angiosperms353 analyses, this script can be run using a filtered mega353 target file as to expand phylogenetic coverage of target file sequences in a custom manner.

### Comparing locus recovery between the default353 target file and the expanded mega353 target file

To compare locus recovery between the default353 versus the expanded mega353 target file we used several datasets, encompassing orders (Asparagales, Sapindales), families (Ericaceae), and genera (*Azorella* Lam., Apiaceae; *Nepenthes* L., Nepenthaceae; *Cyperus* L., Cyperaceae (Larridon et al., 2019); *Bulbophyllum* Thouars., Orchidaceae), as well as the dataset used to test the bait kit in the original Angiosperms353 publication (i.e. the exemplar Angiosperms353 dataset; Johnson et al., 2019) (Table 1). A target file corresponding to each dataset was produced by filtering the mega353 target file to include sequences for the respective family and/or order, depending on the dataset. Because the exemplar Angiosperms353 dataset included a phylogenetically diverse set of angiosperms, the full mega353 target file was used without filtering. The filtered Orchidaceae target file was expanded using a set of *Bulbophyllum* transcriptomes and the BYO_transcriptomes.py script, to create a third more specific target file for the *Bulbophyllum* dataset, in addition to the family and default target files. HybPiper was used to assemble and extract loci sequences, using a nucleotide target file and the flag to call BWA (Li and Durbin 2009) for each dataset, first using the default353 target file as the reference and then the corresponding filtered mega353 target file. For each sample, 16 CPUs and 16 GB of RAM were allocated.

**Table 1:** Summary of recovery statistics produced by HybPiper comparing the default353 target set to the mega353 target set (filtered by family or order). Values represent averages of each dataset for each target file.

| Dataset<br>(number of samples) | Target file | Percentage of<br>reads on target<br>(average) | Number of<br>genes with<br>sequences<br>(average) | Number of genes at<br>75% of target length<br>(average) | Length of<br>concatenated loci<br>(bp, average) |
| --- | --- | --- | --- | --- | --- |
| Angiosperm353<br>exemplar data (43) | default353 | 19.37% | 279.95 | 85.76 | 127796 |
|  | mega353 | 24.07% | 300.20 | 118.24 | 155311 |
|  | mega353 vs default353 % improvement | 24.28% | 7.23% | 37.88% | 21.53% |
| Asparagales (8) | default353 | 1.40% | 192.20 | 30.60 | 73432 |
|  | Order (Asparagales) | 1.90% | 204.40 | 37.60 | 83959 |
|  | Order vs default353 % improvement | 35.71% | 6.35% | 22.88% | 14.33% |
| *Azorella* (5) | default353 | 15.18% | 287.40 | 76.20 | 122667 |
|  | Family (Apiaceae) | 15.74% | 294.40 | 91.40 | 133728 |
|  | Order (Apiales) | 18.58% | 304.20 | 103.40 | 145317 |
|  | Family vs default353 % improvement | 3.69% | 2.44% | 19.95% | 9.02% |
|  | Order vs default353 % improvement | 22.40% | 5.85% | 35.70% | 18.47% |
| *Bulbophyllum* (12) | default353 | 12.30% | 238.83 | 46.00 | 93043 |
|  | Family (Orchidaceae) | 14.71% | 269.17 | 74.92 | 121924 |
|  | Family + genus<br>(Orchidaceae+*Bulbophyllum*) | 15.08% | 272.83 | 84.25 | 130310 |
|  | Family vs default353 % improvement | 19.58% | 12.70% | 62.86% | 31.04% |
|  | Family+genus vs default353 %<br>improvement | 22.63% | 14.24% | 83.15% | 40.05% |
| Cyperaceae (6) | default353 | 10.30% | 192.17 | 64.17 | 86405 |
|  | Family (Cyperaceae) | 12.00% | 242.50 | 94.33 | 121491 |
|  | Order (Poales) | 13.08% | 244.50 | 92.33 | 123825 |
|  | Family vs default353 % improvement | 16.50% | 26.19% | 47.01% | 40.61% |
|  | Order vs default353 % improvement | 27.02% | 27.23% | 43.90% | 43.31% |
| Ericaceae (12) | default353 | 7.57% | 306.25 | 92.33 | 141518 |
|  | Family (Ericaceae) | 12.05% | 335.67 | 175.25 | 191891 |
|  | Order (Ericales) | 13.05% | 337.75 | 181.00 | 198315 |
|  | Family vs default353 % improvement | 59.25% | 9.61% | 89.80% | 35.59% |
|  | Order vs default353 % improvement | 72.47% | 10.29% | 96.03% | 40.13% |
| *Nepenthes* (8) | default353 | 9.28% | 299.50 | 97.13 | 138639 |
|  | Order (Caryophyllales) | 12.50% | 317.63 | 144.00 | 173801 |
|  | Order vs default353 % improvement | 34.77% | 6.05% | 48.26% | 25.32% |
| Sapindales (6) | default353 | 21.02% | 327.67 | 76.67 | 141497 |
|  | Order (Sapindales) | 28.07% | 342.50 | 195.67 | 203309 |
|  | Order vs default353 % improvement | 33.54% | 4.53% | 155.22% | 43.68% |
|  | Average percentage improvement | 31.0% | 11.1% | 61.9% | 30.3% |
|  | Minimum percentage improvement | 3.7% | 2.4% | 19.9% | 9.0% |
|  | Maximum percentage improvement | 72.5% | 27.2% | 155.2% | 43.7% |

The default353 and filtered mega353 target file results were compared using statistics provided by the HybPiper scripts hybpiper_stats.py and get_seq_lengths.py, averaged across all samples for each dataset (Table 1). Four statistics were considered: 1) percentage of reads on target, i.e. the number of reads for a sample that map to the loci in the target file, 2) number of genes with sequences, or the total number of genes that are in the final locus set for each sample, 3) the number of loci ≥75% of target length, i.e. of those loci in the final dataset, the number that are ≥75% length of the target sequence for that gene, and 4) the concatenated length (bp) of the final loci set for each sample.

For each dataset, the new filtered mega353 target file improved each of these measures (Table 1). The average percentage of reads on target improved by 31% across all datasets (between 3.7% and 72.5%). This had the downstream impact of increasing the number of genes with sequences by an average of 11.1% (20 genes) across all datasets (between 2.4% or seven genes, and 27.2% or 50 genes). A greater increase was found in the number of genes at ≥75% target length, with an average increase of 61.9% (46 genes) across all datasets (between 19.9% or 15 genes, and 155.2% or 119 genes). The total length of the concatenated loci increased by an average of 30% (from an average of 115 kb to an average of 148 kb).

For the *Bulbophyllum* dataset, analyses using the target file with sequences from 12 additional *Bulbophyllum* transcriptomes showed improvements over the filtered Orchidaceae target file, with a 2.5% increase in mapped reads, a 12% increase in genes over 75%, and a 7% increase in concatenated loci length (Table 1).

The first script in the HybPiper pipeline is ‘reads_first.py’, which includes mapping of sequence reads to target references and subsequent assembly, and is the most computationally time-consuming step of the pipeline. For most datasets, using a filtered mega353 target file resulted in a small increase in the number of CPU hours taken by each HybPiper run, because as more reference targets are added the time taken for ‘reads_first.py’ increases (Supplementary Table 2). However, the CPU hours used by HybPiper to run the Angiosperms353 exemplar dataset more than tripled with the mega353 target file compared to the default353 target file. This is because the unfiltered mega353 target file was used to account for the phylogenetic breadth in the dataset, and so each locus was represented by 280 sequences (on average) against which reads were mapped. For this reason, we recommend strategically selecting the phylogenetic rank used to filter the target file (i.e. clade, order or family should be preferred where possible), rather than using the complete mega353 target file. Filtering can be applied using multiple phylogenetic ranks as listed in the control file (see https://github.com/chrisjackson-pellicle/NewTargets) For example, for Malvales, a filtered mega353 target file could comprise the target sequences from the order, in addition to selected outgroup sequences (e.g. Brassicaceae), and a specific 1KP sample name (e.g. UPZX, *Cleome gynandra* L., Cleomaceae).

### Expanding phylogenetic density of target files for custom bait kits with BYO_transcriptomes.py

The input required for the script ‘BYO_transcriptomes.py’ is a target file and a directory of transcriptomes and/or nucleotide sequences corresponding to protein-coding genes and can therefore be used to expand target files from other bait kits. To test this functionality, BYO_transcriptomes.py was used to expand target files for an Asteraceae-specific bait kit (Mandel et al., 2014), and a Hibisceae-specific bait kit (McLay et al. in prep.).

The Asteraceae bait kit was designed using *Helianthus annuus* L. (sunflower; Asteroideae), *Lactuca sativa* L. (lettuce; Cichorioideae), and *Carthamus tinctorius* L. (safflower; Carduoideae). The Asteraceae target file (comprising only the *H. annuus* and *L. sativa* target sequences) was expanded using 1KP transcriptomes selected as they were closely related to Asteraceae tribe Gnaphalieae (Supplementary Table 3). The Hibisceae-specific bait kit was designed using three Hibisceae transcriptomes, *Abelmoschus esculentus* (L.) Moench, *Hibiscus cannibinus* L., and *Hibiscus syriacus* L.. The Hibisceae target file was expanded using available sequence data from the other Malvaceae subfamily Malvoideae tribes, Malveae and Gossypieae (Supplementary Table 3).

Each new target file was compared to its default target file using HybPiper with the approach described above. Seven representative samples from Asteraceae tribe Gnaphalieae, captured using the Asteraceae bait kit (Mandel et al., 2014) were used to compare the default Asteraceae target file (two targets per locus) to the expanded Asteraceae target file (average of 3.88 targets per locus). Five representative taxa from Malvaceae tribes Malveae and Gossypieae, captured using the Hibisceae bait kit, were used to compare the default Hibisceae target file (average of 2.5 targets per locus) to the expanded Malvoideae target file (average 4.34 targets per locus). Locus recovery was improved using the expanded target file for both datasets. This improvement was more pronounced with the expanded Asteraceae target file, with a 31% increase in the number of genes at ≥75% of the target length, and a 22% increase in concatenated loci length (Supplementary Table 4).

## CONCLUSION

We have demonstrated that sequence recovery for a universal sequence capture bait kit can be substantially improved by appropriate tailoring of target files to the group under study. To enable the best possible locus recovery from Angiosperms353 capture data, we have developed an expanded target file using 1KP transcriptomes. As the Angiosperms353 bait kit is becoming increasingly widely used, tools such as we have developed here will allow researchers to optimise use of their target enrichment sequence data by assembling more and longer loci, increasing cost efficiency, dataset combinability, and likely enabling better phylogenetic outcomes. Furthermore, our BYO_transcriptomes.py script can be used to incorporate additional target sequences from any available transcriptome, and we have shown that this tool can be used to improve locus recovery using target files other than the Angiosperms353 bait kit. With the growing number of transcriptomes and whole genome data becoming available in public repositories, the approach developed here will prove to be an increasingly valuable resource for efficient recovery of target enrichment data.

## ACKNOWLEDGEMENTS

WJB, FF and EMJ were supported by grants from the Calleva Foundation and the Sackler Trust to the Plant and Fungal Tree of Life Project (PAFTOL) at the Royal Botanic Gardens, Kew. Matt Johnson is thanked for providing useful feedback in the early stages of this project. JLB and BFG were supported by a Herman Slade Foundation grant (HSF1608).

---

**Supplementary Figure 1:**
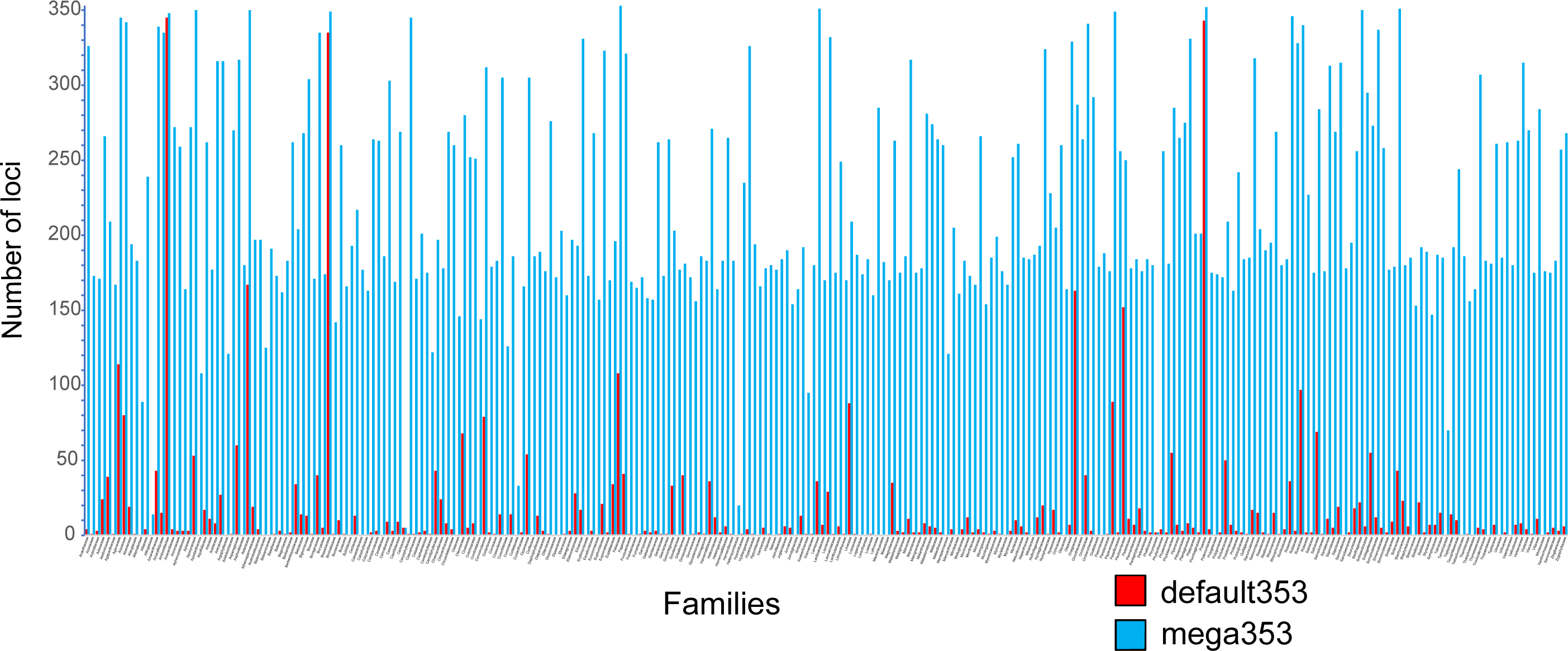
Comparing the number of genes represented for each family in the default353 (red) or mega353 (blue) target files.

**Supplementary Table 1:** The number of target sequences in the default353 target file compared to the mega353 target file, including the average number of targets per locus, and the average number of orders and families for each locus.

|  | **default353** | **mega353** |
| --- | --- | --- |
| Total number of target sequences | 4780 | 98994 |
| Average number of target reference sequences per locus<br>(minimum – maximum) | 13.5<br>(6–18) | 280<br>(17–373) |
| Average number of orders for each locus<br>(minimum – maximum, total) | 13.5<br>(6–18, 55) | 49.8<br>(13–57, 57) |
| Average number of families for each locus<br>(minimum – maximum, total) | 13.5<br>(6–18, 226) | 170<br>(14–214, 276) |

**Supplementary Table 2:** CPU hours used by the HybPiper pipeline to complete for each dataset and each target file. HybPiper was allocated 16 CPUs and 16 GB of RAM for each dataset

| **Dataset** | **Target file** | **CPU hours** |
| --- | --- | --- |
| Angiosperm353 exemplar data | default353 | 108 |
|  | mega353 | 363.2 |
| Asparagales | default353 | 9.6 |
|  | Order | 11 |
| *Azorella* | default353 | 97.6 |
|  | Family | 100.2 |
|  | Order | 105.6 |
| Cyperaceae | default353 | 22.2 |
|  | Family | 21.2 |
|  | Order | 21.6 |
| Ericaceae | default353 | 176.2 |
|  | Family | 282.3 |
|  | Order | 264.2 |
| Nepenthes | default353 | 42 |
|  | Order | 42.9 |
| Sapindales | default353 | 54.9 |
|  | Order | 66.4 |

**Supplementary Table 3:** Samples used to expand the custom bait kit target files using ‘BYO_transcriptomes.py’

| Bait kit dataset | Samples added (source) |
| --- | --- |
| Asteraceae –<br>Mandel et al., 2014 | *Senecio rowleyanus* H.Jacobsen – 1KP BMSE<br>*Leontopodium alpinum* (Ten.) A.Huet ex Hand.-Mazz.– 1KP DOVJ<br>*Matricaria matricarioides* (Less.) Porter – 1KP OAGK<br>*Solidago canadensis* L. – 1KP TEZA |
| Hibisceae – McLay<br>et al.in prep | *Hoheria angustifolia* Raoul – 1KP ZSAB<br>*Gossypium australe* F.Muell – GenBank PRJNA513946 |

**Supplementary Table 4:** Comparing custom bait kit target files (Asteracae/Hibisceae) that were expanded using BYO_transcriptomes.py. Values represent averages of each dataset for each target file.

| Bait kit dataset<br>(number of<br>samples tested) | Target file | Percentage of<br>reads on target<br>(average) | Number of genes<br>with sequences<br>(average) | Number of genes at<br>75% of target<br>length<br>(average) | Length of<br>concatenated loci<br>(bp, average) |
| --- | --- | --- | --- | --- | --- |
| Asteraceae –<br>Mandel et al. 2014<br>(7) | Asteraceae<br>target file | 15.00% | 560.17 | 332.67 | 158938.5 |
|  | Expanded<br>target file | 23.00% | 665.5 | 435.83 | 194377 |
|  | %<br>improvement | 54.30% | 18.80% | 31.01% | 22.30% |
| Hibisceae – McLay<br>et al.in prep<br>(5) | Hibisceae<br>target file | 24.78% | 504.8 | 449.6 | 247443.6 |
|  | Expanded<br>target file | 27.84% | 512.6 | 461.6 | 267151.8 |
|  | %<br>improvement | 12.35% | 1.55% | 2.67% | 7.96% |

